# NEMO- and RelA-dependent NF-κB Signaling Promotes Small Cell Lung Cancer

**DOI:** 10.1101/2022.07.29.501988

**Authors:** Lioba Koerner, Tsun-Po Yang, Marcel Schmiel, Martin Peifer, Reinhard Buettner, Manolis Pasparakis

## Abstract

Small cell lung cancer (SCLC) is an aggressive type of lung cancer driven by combined loss of the tumor suppressors *RB1* and *TP53*. SCLC is highly metastatic and despite good initial response to chemotherapy patients usually relapse, resulting in poor survival. Therefore, better understanding of the mechanisms driving SCLC pathogenesis is required to identify new therapeutic targets. Here we identified a critical role of the IKK/NF-κB signaling pathway in SCLC development. Using a relevant mouse model of SCLC, we found that ablation of NEMO/IKKγ, the regulatory subunit of the IKK complex that is essential for activation of canonical NF-κB signaling, strongly delayed the onset and growth of SCLC resulting in considerably prolonged survival. In addition, ablation of the main NF-κB family member p65/RelA also delayed the onset and growth of SCLC and prolonged survival, albeit to a lesser extent than NEMO. Interestingly, constitutive activation of IKK/NF-κB signaling within the tumor cells did not exacerbate the pathogenesis of SCLC, suggesting that endogenous NF-κB levels are sufficient to fully support tumor development. Moreover, TNFR1 deficiency did not affect the development of SCLC, showing that TNF signaling does not play an important role in this tumor type. Taken together, our results revealed that IKK/NF-κB signaling plays an important role in promoting SCLC, identifying the IKK/NF-κB pathway as a promising therapeutic target.

## Introduction

Small cell lung cancer (SCLC) is a very aggressive lung cancer subtype with a median survival of ~1 year for patients with metastatic disease ^1^. SCLC is characterized by early metastatic spread, neuroendocrine differentiation and small tumor cells ^1, 2, 3, 4^. Patients often initially respond to chemotherapy but in most cases the tumors subsequently relapse and become resistant to cytotoxic treatments ^1, 5^. Comprehensive genomic profiling of SCLC revealed a bi-allelic loss of *RB1* and *TP53* in all patients, showing that loss of these two tumor suppressors is obligatory in SCLC ^6, 7, 8, 9, 10^. However, the mechanisms determining the aggressive nature of SCLC and its increased metastatic potential remain poorly understood ^1^. Therefore, a better understanding of the molecular pathways that determine SCLC initiation, progression and metastasis will be critical to develop new therapeutic approaches.

The interplay between tumor and immune cells plays an essential role in cancer progression and critically determines tumor development. Indeed, inflammation has been recognized as one of the hallmarks of cancer ^11^. While chronic inflammation can fuel tumor growth, an adaptive immune response against tumor-specific antigens can also trigger anti-cancer immunity, resulting in elimination of transformed cells ^11^. SCLC ranks among the tumor entities with the highest tumor mutational burden, likely due to it cigarette smoke-induced nature, which is expected to result in large number of neoantigens. However, immunotherapies based on checkpoint inhibitors had limited success with only a small percentage of patients responding ^1^. Tumor necrosis factor (TNF) is a potent pro-inflammatory cytokine exerting both tumor-promoting and anti-cancer effects ^12^. TNF was discovered as a factor inducing the death of tumor cells but was subsequently shown to promote tumor development in different models, such as DMBA/TPA induced skin carcinogenesis and obesity-associated liver cancer ^13, 14^. TNF signaling via TNFR1 regulates inflammation, cell survival and death by inducing distinct intracellular signaling cascades ^15^. TNFR1 stimulation induces the activation of the inhibitor of NF-κB (IκB) kinase (IKK) complex, resulting in the nuclear accumulation of NF-κB promoting the transcription of genes regulating inflammation and cell survival ^15, 16^. The IKK complex consists of the regulatory subunit NEMO (also termed IKKγ) and the catalytic subunits IKK1 and IKK2 (also termed IKKa and IKKß, respectively ^16, 17, 18^. NEMO is essential for activation of canonical NF-κB signaling, which primarily depends on IKK2 kinase activity and the RelA NF-κB subunit ^16, 17, 18^. IKK/NF-κB signaling has emerged as a crucial driver of tumor growth and progression ^17, 19^. Multiple studies in mouse models demonstrated that NF-κB signaling promotes tumorigenesis in several cancer entities, including colitis-associated colon cancer ^20^, mammary tumors ^21, 22, 23^, DMBA/TPA-induced skin carcinogenesis ^24^, as well as *Kras* mutation-driven lung adenocarcinoma ^25, 26, 27^. However, NF-κB signaling has also been shown to exhibit tumor suppressing functions in different tissues and models of carcinogenesis. Inhibition of NF-κB signaling in human keratinocytes promoted Ras-mediated oncogenic transformation in a xenograft model ^28^ and NF-κB inhibition via expression of a dominant-negative mutant IκBα super-repressor (IκBαSR) in murine skin triggered the development of squamous cell carcinoma ^29, 30^. Additionally, liver parenchymal cell-specific knockout of NEMO caused the spontaneous development of chronic hepatitis and hepatocellular carcinoma (HCC) in mice ^31, 32^, whereas ablation of IKK2 in hepatocytes led to increased diethylnitrosamine (DEN)-induced liver tumorigenesis ^33^. Interestingly, NF-κB activation has been proposed to induce T-cell mediated immune surveillance and therefore tumor rejection in lung adenocarcinoma ^34^. Thus, the role of NF-κB signaling has been extensively studied in various tumor entities, amongst them *Kras*-driven lung adenocarcinoma, however, its function in SCLC remains elusive.

Here we aimed to study the role of TNFR1 and NF-κB signaling in SCLC using an autochthonous model of SCLC induced by the simultaneous ablation of *Rb1* and *Tp53* in mouse lung epithelial cells. We show that inhibition of NF-κB signaling by depletion of NEMO or RelA delayed tumor onset, slowed tumor growth and prolonged mouse survival. Surprisingly, neither constitutive activation of NF-κB signaling, nor ablation of TNFR1 resulted in changes in SCLC development or progression. Together, these results show that NEMO and RelA-dependent NF-kB signaling plays a critical role in SCLC.

## Materials and Methods

### Mice

*Rb*^FL/FL 35^, *Tp53*^FL/FL 36^, *Nemo*^FL/FL 37^, *Rela*^FL/FL 38^, *R26*^*LSL.IKK2ca* 39^, *Tnfr1*^FL/FL 40^ and *Tnfr1^-/-^*^41^ mice have been described before. Mice used in these experiments were kept on a mixed C57Bl6/J/N background. Before MR imaging, mice were maintained at the specific pathogen-free animal facility of the CECAD Research Center, University of Cologne, under a 12-hour dark/12-hour light cycle in individually ventilated cages (Greenline GM500; Tecniplast) at 22 (±2) °C and a relative humidity of 55 (±5) %. They were given a sterilized commercial pelleted diet (Ssniff Spezialdiäten GmbH) as well as water ad libitum. All animal procedures were conducted in accordance with European, national and institutional guidelines and experimental protocols were approved by local government authorities (Landesamt für Natur, Umwelt und Verbraucherschutz Nordrhein-Westfalen). Animals requiring medical attention were provided with appropriate care and excluded from the studies described when reaching pre-determined criteria of disease severity. No other exclusion criteria existed. Mouse studies as well as immunohistochemical assessment of pathology and evaluation of MR imaging were performed in a blinded fashion.

### RP mouse model & MRI scans

For induction of lung tumor formation, 8–12-week-old mice were anesthetized with Ketavet (100mg/kg)/Rompun (20mg/kg) by intraperitoneal injection, followed by intratracheal application of replication-deficient Cre-expressing adenovirus (Ad5-CMV-Cre, 2.5 x 10^7^ PFU, University of Iowa). Mice were inhaled in three different cohorts, every cohort including all genotypes used in this study. Only cohort 1 did not include *Rb1*^FL/FL^*Tp53*^FL/FL^*Nemo*^FL/FL^ mice. One *Rb1*^FL/FL^*Tp53*^FL/FL^*Tnfr1^-/-^* mouse was inhaled at a later timepoint. All experiments carried out were pooled. Starting 20 weeks after AdenoCre inhalation, tumor development was monitored bi-weekly by magnetic resonance imaging by using the MRI (A 3.0 T Philips Achieva clinical MRI) in combination with a specific mouse solenoid coil (Philips Hamburg, Germany). MR images were acquired using turbo-spin echo (TSE) sequence (repetition time = 3819ms, echo time = 60ms, field of view = 40 x 40 x 20 mm^3^, reconstructed voxel size = 0.13 x 0.13 x 1.0 mm^3^) under 2,5% isoflurane anesthesia. Resulting MR images were analyzed blindly by marking regions of interests employing Horos software. For MRI analysis, mice were maintained in the animal facility of the nuclear medicine, University Hospital Cologne, in individually ventilated cages at 12h/12h light/dark cycle, 55 (±10%) humidity and 22 (±2) °C. All animal experiments were approved by local government authorities (Landesamt für Natur, Umwelt und Verbraucherschutz, Nordrhein-Westfalen, Germany). All animal experiments were conducted in compliance with european, national and institutional guidelines on animal welfare. Animals requiring medical attention were provided with appropriate care and excluded from the studies described when reaching pre-determined criteria of disease severity. No other exclusion criteria existed.

### Tissue preparation

Mice were sacrificed using cervical dislocation. For histopathological analysis the trachea was injected with 4% PFA to inflate and the lung. Lung tissue was fixed in 4% PFA O/N at 4°C. Small pieces of tumor tissues were snap frozen on dry ice for RNA and protein expression analysis and stored at −80°C until further processing. Tumors were cut from the lungs, in order to isolate cell lines as described below.

### Cell culture

Tumors were isolated from RP-mice at the humane endpoint. The tumors were incubated in 10X T rypLE™ (ThermoFisher Scientific #A1217701) at 37°C and 5% CO_2_ for 20 minutes. Roswell Park Memorial Institute (RPMI) 1640 medium containing 10% FCS and 1% P/S was added and the tissue was incubated at 37°C, 5% CO_2_ overnight. After incubation, remaining tumor tissue was removed from the culture and cells were grown at 37°C, 5% CO_2_, washed with PBS every third day and supplied with new RPMI medium + 1%P/S + 10% FCS until they grew confluent. Cell lines were then maintained at 37°C and 5% CO_2_.

### Histologic analysis

Formalin-fixed paraffin embedded (FFPE) 4 μm-thick lung tissue sections were de-paraffinized using xylene and re-hydrated with decreasing ethanol concentrations (100%, 96%, 75%, 0%). The tissue sections were stained in haematoxylin for 2 minutes, 15 minutes differentiated in tap water and stained for 1 minute with eosin. Then, sections were de-hydrated using increasing ethanol concentrations and fixed in xylene. The slides were mounted in Entellan. In addition, FFPE lungs were stained for Ki-67 (Cell Marque 275-R10), CD45 (BD 550539) and CD56 (Zytomed RBK050).

### Immunoblotting

For immunoblot analyses, 3 x 10^5^ cells were seeded in 6-well plates and cultured O/N. Cells were lysed in RIPA buffer (HEPES 20 nM, NaCl 350 mM, MgCl2 1mM, EDTA 0.5mM, EGTA 0.1mM, Glycerol 20%, Nonident P-40 1%) supplemented with protease and phosphatase inhibitor tablets (Roche) for 20 minutes on ice. Proteins from tumor tissue were isolated using precellys 24 tissue homogenizer (bertin instruments). Protein concentration was measured using PIERCE 660nm Protein Assay Reagent (Thermo Scientific, #22660) and BSA standard. Lysate concentration was adjusted to 5 μg/μl and 2x Laemmli sample buffer (Bio-Rad 1610737) was added. Samples were boiled at 95°C for 8 minutes.

Cell lysates were separated using Sodium dodecyl-sulfate polyacrylamide gel electrophoresis and transferred to polyvinylidene difluoride membranes (IPVH00010, Millipore) at 80V for 3h at 4°C. Membranes were blocked using 5% milk in 0.1% PBST for 1 h, washed three times with 0.1% PBST and probed with following primary antibodies: α-NEMO (homemade 1:1000), a-p65 (Cell Signaling 3179 1:1000), a-IKK2 (Cell Signaling 2684S 1:1000) and a-Tubulin (Sigma T6074 1:1000) O/N at 4°C. Membranes were washed three times with 0.1% PBS-T and were incubated with secondary horseradish peroxidase-coupled antibodies for 1h at RT (GE Healthcare, Jackson ImmunoResearch, 1:10000). ECL Western Blotting Detection Reagent (RPN2106, GE Healthcare) was used to detect the proteins. Membranes were stripped if necessary, using stripping buffer (ThermoScientific, 21059) for 15 minutes at RT.

### 3’ mRNA sequencing analysis

RNA isolation from tumor tissue was performed using a NucleoSpin RNA isolation kit (Macherey Nagel Ref. 740955.250).

RNA quality was evaluated based on OD260/280 and OD260/230 ratios as well as on RNA integrity number (RIN). For determination of gene expression, the Quant 3’mRNAseq Library Prep Kit FWD for Illumina (Lexogen). Samples with RIN < 4, OD260/260 <1.8 or OD 260/230 < 1.5 were excluded from the analysis. Five mice per genotype were used. Single-end sequencing reads were aligned to Ensembl GRCm38 (mm10) cDNA sequences using kallisto 0.43.1 ^42^ with default average fragment length parameters. Transcript-level transcripts per million (TPM) normalisation were estimated, and gene-level aggregated TPMs were calculated using sleuth 0.29.0 ^43^resulting in 35,930 genes with a Ensembl BioMart annotation. Only 18,383 of those were considered expressed (median TPM > 0) across 41 transcriptomes and subsequently used for further analysis.

### Gene set enrichment analysis (GSEA)

All expressed genes were first weighted using the calculation shown below to convert the two-dimension dynamics (i.e., fold changes and p-value significances) derived from differential analysis into a one-dimension gene list. Then this pre-ranked gene list was used to run against the hallmark gene sets downloaded from MSigDB mouse draft database 0.3 to test for enrichment using GSEA tool 4.2.3 ^44^.

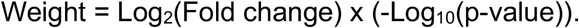

### Statistical analysis

Data shown in graphs display mean. Wilcoxon rank-sum/ Mann-Whitney *U* test was used to test for differential gene expression between two non-parametric groups, and the results were visualised via volcano plots using purpose-written R scripts.For comparison of more than two groups, Kruskal-Wallis Test was used. The Log-Rank (Mantel-Cox) test was used in order to compare survival curves and tumor onset of mice. *p ≤ 0.05; **p ≤ 0.01; ***p ≤ 0.005, ****p ≤ 0.001 for all figures. All statistical analysis was performed with Prism6, GraphPad. No data were excluded.

### Data availability statement

The RNA-seq data reported in the manuscript have been deposited in the European Nucleotide Archive (ENA) at EMBL-EBI under accession number PRJEB53995 (https://www.ebi.ac.uk/ena/browser/view/PRJEB53995).

## Results

### Critical role of NEMO in SCLC development

To study the role of NF-kB signaling in SCLC, we employed a well-characterized genetically engineered mouse model of the disease based on combined ablation of *Rb1* and *Tp53* in mouse lung epithelial cells via adenovirus-mediated delivery of Cre recombinase, which causes the development of SCLC within 9 months (Figure 1A). This mouse model recapitulates the key features of human SCLC, including the histopathological and immunological phenotype ^45^. To inhibit canonical NF-kB signaling, we chose to target NEMO, the regulatory subunit of the IKK complex that is essential for canonical NF-kB activation, and p65/RelA, which is the NF-kB subunit primarily responsible for the transcriptional activation of canonical NF-kB target genes ^16, 18^. To address the role of NEMO in SCLC we crossed mice carrying loxP-flanked *Nemo* alleles to mice carrying loxP-flanked *Rb1* and *Tp53* alleles. At the age of 8-12 weeks, *Rb1*^FL/FL^ *Tp53*^FL/FL^ as well as *Rb1*^FL/FL^ *Tp53*^FL/FL^ *Nemo^FL/FL^* mice were inhaled with adenovirus expressing Cre recombinase (Ad-Cre), which results in Cre-mediated deletion of the respective loxP-flanked alleles in lung epithelial cells. Starting at 20 weeks after Ad-Cre inhalation, mice were monitored for tumor development by magnetic resonance imaging (MRI) scanning biweekly (Figure 1A, B). Ad-Cre-inhaled *Rb1*^FL/FL^ *Tp53*^FL/FL^ mice developed tumor lesions with a median tumor onset of 24 weeks after inhalation (Figure 1B, C). In contrast, *Rb1*^FL/FL^ *Tp53*^FL/FL^ *Nemo^FL/FL^* mice showed a strong delay in tumor development with a median tumor onset of 37 weeks after Ad-Cre inhalation (Figure 1B, C). Overall, 10 out of 11 *Rb1*^FL/FL^ *Tp53*^FL/FL^ *Nemo^FL/FL^* mice eventually developed tumors, whereas one mouse did not show signs of tumor development as late as 400 days after Ad-Cre inhalation (Figure 1B, C). At 26 weeks after Ad-Cre inhalation, we detected the presence of several lung tumors in 8 out of 17 *Rb1*^FL/FL^ *Tp53*^FL/FL^ mice, whereas none of the *Rb1*^FL/FL^ *Tp53*^FL/FL^ *Nemo^FL/FL^* mice showed lung tumor development at this stage (Figure 1D). Longitudinal measurements of tumor volume by MRI showed that, in addition to the delayed onset, lung tumors in *Rb1*^FL/FL^ *Tp53*^FL/FL^ *Nemo^FL/FL^* mice displayed considerably reduced growth compared to *Rb1*^FL/FL^ *Tp53*^FL/FL^ mice during the first six weeks after tumor detection (Figure 1E). In addition to biweekly MRI scans, all mice were screened regularly for distress symptoms and were sacrificed humanely when reaching pre-determined termination criteria. In line with delayed tumor onset and reduced tumor growth, *Rb1*^FL/FL^ *Tp53*^FL/FL^ *Nemo^FL/FL^* mice showed a significantly increased overall survival with a median survival of 358 days, compared to *Rb1*^FL/FL^ *Tp53*^FL/FL^ mice that had a median survival of 247 days (Figure 1F). Collectively, ablation of NEMO strongly delayed tumor onset, reduced tumor growth and significantly prolonged survival in a relevant mouse model of SCLC induced by combined inactivation of RB1 and TP53.

**Figure 1.**
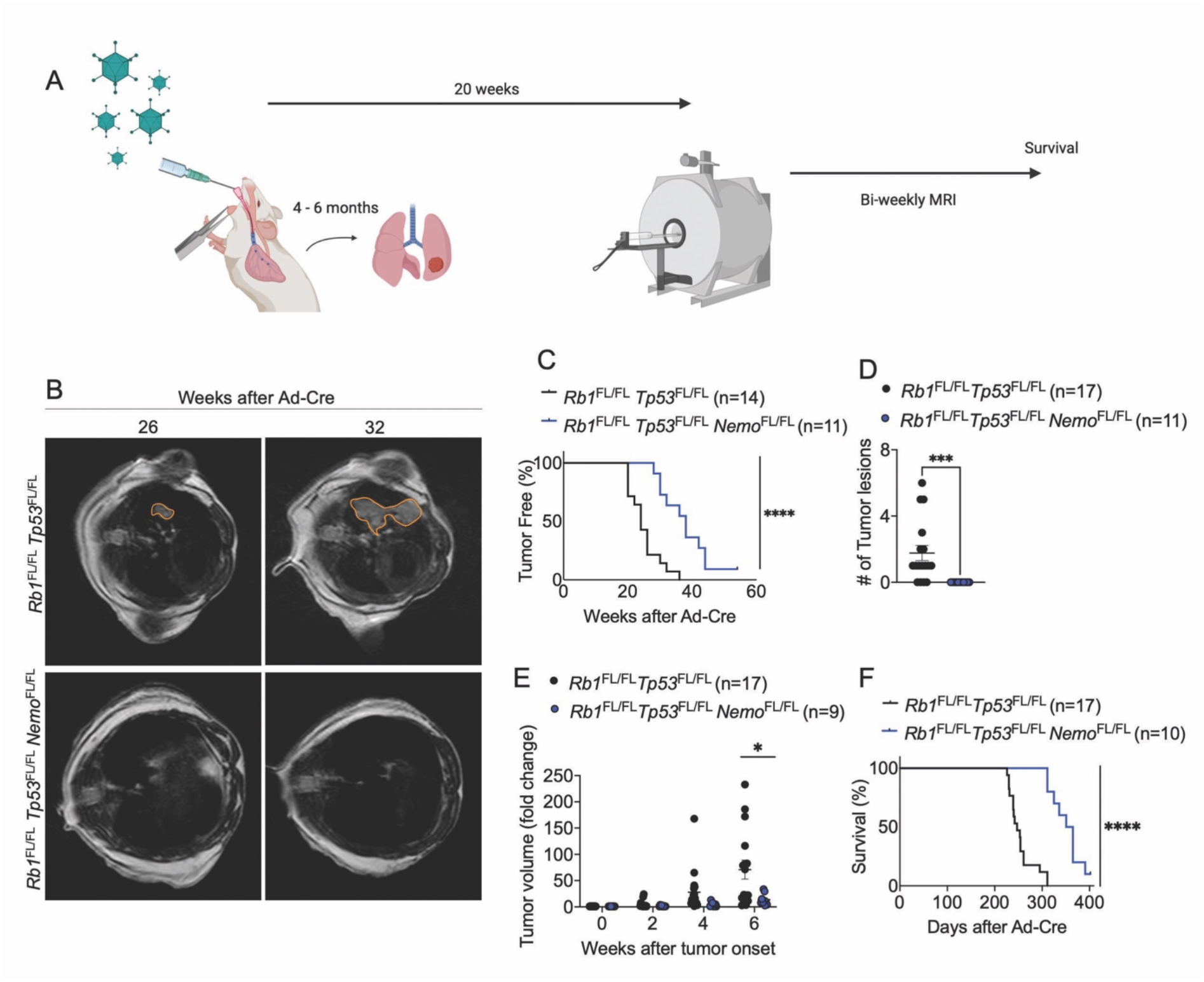
Epithelial NEMO ablation delays tumor onset and prolongs survival in a mouse model of SCLC. **(A)** Schematic showing intratracheal Ad-Cre inhalation and experimental procedure for the generation and analysis of a mouse model of SCLC (created with BioRender.com) **(B)** Representative images of MRI scans of *Rb1*^FL/FL^ *Tp53*^FL/FL^ (n=17) and *Rb1*^FL/FL^ *Tp53*^FL/FL^ *Nemo*^FL/FL^ (n=9) mice 26 and 32 weeks after Ad-Cre inhalation. Tumor areas were marked using Horos Software. **(C)** Graph depicting tumor onset assessed via MR imaging of mice with indicated genotypes. ****p < 0.0001, Log-rank test. **(D)** Graph depicting the number of tumor lesions 6 weeks after Ad-Cre inhalation of the indicated genotypes. ***p = 0.001, Mann-Whitney Test. Mean ± SEM. Each dot represents one mouse. **(E)** Graph depicting tumor fold change assessed via MR imaging 2-6 weeks after Ad-Cre inhalation for indicated genotypes. Each dot represents one mouse. Bars represent mean ± SEM. *p < 0.05, Mann-Whitney Test. **(F)** Graph depicting survival of mice with indicated genotypes. ****p < 0.0001, Log-rank test.

Despite the delayed onset and reduced growth of tumors in Ad-Cre-treated *Rb1*^FL/FL^ *Tp53*^FL/FL^ *Nemo*^FL/FL^ compared to *Rb1*^FL/FL^ *Tp53*^FL/FL^ mice, macroscopic examination of lungs dissected from mice sacrificed at the humane endpoint revealed similar tumor burden in both genotypes (Figure 2A). In addition to the lung tumors, SCLC metastasis to the liver was also detected in about half of the mice with no difference in metastatic prevalence between the genotypes (Figure 2B). Histopathological analysis of lung sections showed the presence of tumors displaying classical characteristics of SCLC, such as homogenous tissue composed out of small tumor cells and expression of CD56, a marker frequently used in SCLC diagnosis ^46^ (Figure 2C). Immunohistochemical staining with antibodies against Ki-67 revealed similar numbers of proliferating cells in tumors from *Rb1*^FL/FL^ *Tp53*^FL/FL^ *Nemo*^FL/FL^ compared to *Rb1*^FL/FL^ *Tp53*^FL/FL^ mice (Figure 2C). Considering that NF-κB signaling regulates various aspects of innate and adaptive immunity ^47^, we additionally investigated tumor immune cell infiltration by immunostaining for CD45, a marker expressed in all immune cells. Immunostaining for CD45 revealed the presence of immune cells in lung tissue surrounding the malignant lesions, however, we did not observe prominent immune cell infiltration within the tumor mass in either *Rb1*^FL/FL^ *Tp53*^FL/FL^ *Nemo*^FL/FL^ or *Rb1*^FL/FL^ *Tp53*^FL/FL^ mice (Figure 2D). To obtain insights into possible effects of NEMO deficiency in the transcriptional profile of SCLC, we performed RNA sequencing (RNAseq) analysis in RNA isolated from lung tumors dissected from *Rb1*^FL/FL^ *Tp53*^FL/FL^ *Nemo*^FL/FL^ and *Rb1*^FL/FL^ *Tp53*^FL/FL^ mice. Surprisingly, we did not find considerable differences in the gene transcriptional profile of the tumors between the two genotypes (Figure 2E). Specifically, we only found only 6 genes significantly upregulated and 6 genes significantly downregulated in tumors from *Rb1*^FL/FL^ *Tp53*^FL/FL^ *Nemo*^FL/FL^ compared to *Rb1*^FL/FL^ *Tp53*^FL/FL^ mice (Figure F). Therefore, analysis of tumors at the time of humane sacrifice did not reveal differences in tumor size, proliferation or immune cell infiltration between *Rb1*^FL/FL^ *Tp53*^FL/FL^ *Nemo*^FL/FL^ and *Rb1*^FL/FL^ *Tp53*^FL/FL^ mice.

**Figure 2.**
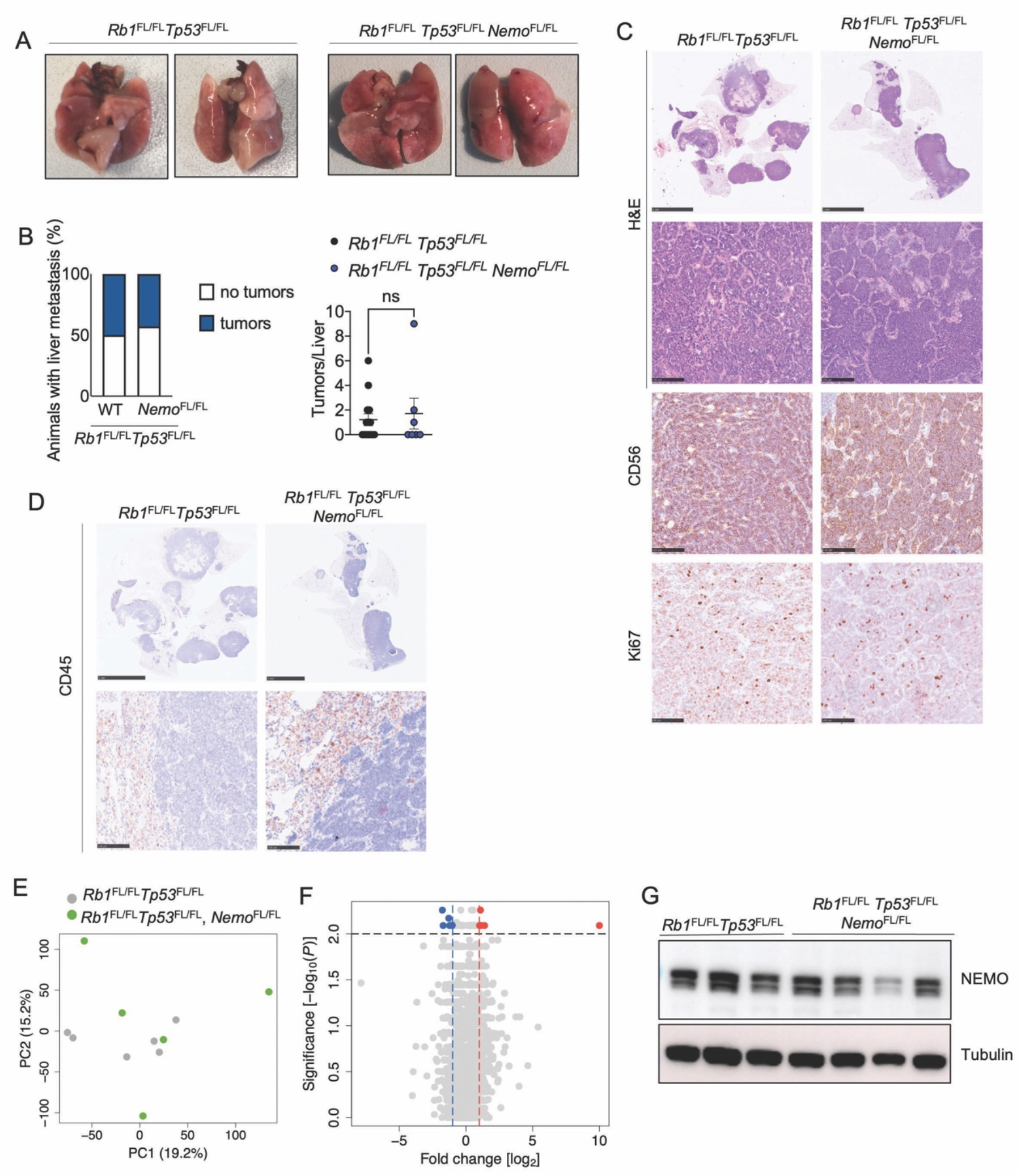
Histological and molecular analysis of lung tumors in *Rb1*^FL/FL^ *Tp53*^FL/FL^ *Nemo*^FL/FL^ mice. **(A)** Representative photographs of lungs of *Rb1*^FL/FL^ *Tp53*^FL/FL^ (n=17) and *Rb1*^FL/FL^ *Tp53*^FL/FL^ *Nemo*^FL/FL^ (n=10) mice sacrificed at the humane endpoint. **(B)** Graphs depicting the percentage of mice with liver metastasis and the amount of liver tumor lesions at the humane endpoint. n= 7 for *Rb1*^FL/FL^ *Tp53*^FL/FL^ *Nemo*^FL/FL^, n=14 for *Rb1*^FL/FL^ *Tp53*^FL/FL^. Mann-Whitney test. Mean ± SEM. **(C)** Representative images of lung sections from mice with indicated genotypes sacrificed at the humane endpoint, stained with H&E or immunostained for CD56 and for Ki67. Scale bars = 5mm & 100μm (CD56: n=8 for *Rb1*^FL/FL^ *Tp53*^FL/FL^ *Nemo*^FL/FL^, n=11 for *Rb1*^FL/FL^ *Tp53*^FL/FL^; Ki67: n=6 for *Rb1*^FL/FL^ *Tp53*^FL/FL^ *Nemo*^FL/FL^, n=13 for *Rb1*^FL/FL^ *Tp53*^FL/FL^). **(D)** Representative images of lung sections from mice at the humane endpoint immunostained for CD45. Scale bars = 5 mm (top) and 100μm (bottom), (n=5 for *Rb1*^FL/FL^ *Tp53*^FL/FL^ *Nemo*^FL/FL^, n=13 for *Rb1*^FL/FL^ *Tp53*^FL/FL^). **(E-F)** Principal component analysis (PCA) (**E**) and Volcano plot (**F**) of RNA seq data from tumor tissues of *Rb1*^FL/FL^ *Tp53*^FL/FL^ *Nemo*^FL/FL^ (n=6) compared to *Rb1*^FL/FL^ *Tp53*^FL/FL^ (n=5) mice. Genes that were found significantly upregulated (p < 0.01, log2(|FC|) > 2) or downregulated (p < 0.01, log2(|FC|) < 2) in tumors from *Rb1*^FL/FL^ *Tp53*^FL/FL^ *Nemo*^FL/FL^ compared to *Rb1*^FL/FL^ *Tp53*^FL/FL^ mice are indicated with red or blue dots, respectively**. (G)** Representative immunoblot analysis of protein extracts from SCLC cell lines isolated from *Rb1*^FL/FL^ *Tp53*^FL/FL^ and *Rb1*^FL/FL^ *Tp53*^FL/FL^ *Nemo*^FL/FL^ mice at the humane endpoint (n=4).

To assess whether tumors developing in Ad-Cre-inhaled *Rb1*^FL/FL^ *Tp53*^FL/FL^ *Nemo*^FL/FL^ mice have lost the expression of NEMO, we isolated SCLC cell lines from the lungs of mice sacrificed at the humane endpoint and analyzed NEMO protein expression by immunoblotting. Surprisingly, we found that NEMO was expressed in all tumor cell lines isolated from *Rb1*^FL/FL^ *Tp53*^FL/FL^ *Nemo*^FL/FL^ mice, with only one out of four cell lines showing reduced levels of NEMO protein compared to cell lines from *Rb1*^FL/FL^ *Tp53*^FL/FL^ mice (Figure 2G). Therefore, tumors developing in *Rb1*^FL/FL^ *Tp53*^FL/FL^ *Nemo*^FL/FL^ mice express NEMO, suggesting that they are derived from cells that recombined the *Rb1*^FL/FL^ and *Tp53*^FL/FL^ alleles, thus losing expression of both tumor suppressors, but failed to undergo recombination of the *Nemo*^FL^ alleles. These findings indicate that tumor development in *Rb1*^FL/FL^ *Tp53*^FL/FL^ *Nemo*^FL/FL^ mice appears to be driven by the selection of clones that have escaped NEMO deletion, which could also contribute to the kinetics observed with a strong delay in tumor onset and progression. Taken together, these results revealed an essential role of NEMO in SCLC.

### Lack of RelA delays tumor onset and prolongs mouse survival in SCLC

NEMO is essential for canonical NF-κB activation but has also important, NF-κB-independent functions in preventing cell death ^31, 48^. Thus, to specifically address the role of NF-κB, we chose to target RelA, the NF-κB subunit that is critical for canonical NF-κB-mediated gene transcription induction ^16, 18^, in addition to NEMO. To this end, we generated *Rb1*^FL/FL^ *Tp53*^FL/FL^ *Rela*^FL/FL^ mice and assessed SCLC development induced by inhalation of Ad-Cre. Assessment of lung tumor presence by MRI revealed that *Rb1*^FL/FL^ *Tp53*^FL/FL^ *Rela*^FL/FL^ mice showed considerably delayed tumor development with a median tumor onset of 28 weeks compared to 24 weeks in *Rb1*^FL/FL^ *Tp53*^FL/FL^ mice (Figure 3A, B). Quantification of tumor lesions at 26 weeks after Ad-Cre inhalation revealed that *Rb1*^FL/FL^ *Tp53*^FL/FL^ *Rela*^FL/FL^ mice showed a trend towards reduced tumor presence compared to *Rb1*^FL/FL^ *Tp53*^FL/FL^ mice, which however did not reach statistical significance (Figure 3C). Moreover, longitudinal measurement of tumor volume by MRI did not reveal statistically significant changes between the two groups, although *Rb1*^FL/FL^ *Tp53*^FL/FL^ *Rela*^FL/FL^ mice showed a trend towards reduced tumor growth compared to *Rb1*^FL/FL^ *Tp53*^FL/FL^ mice (Figure 3D). In line with delayed tumor onset and reduced tumor growth, *Rb1*^FL/FL^ *Tp53*^FL/FL^ *Rela*^FL/FL^ mice showed significantly increased overall survival with a median survival of 311 days compared to *Rb1*^FL/FL^ *Tp53*^FL/FL^ mice that had a median survival of 247 days (Figure 3E).

**Figure 3.**
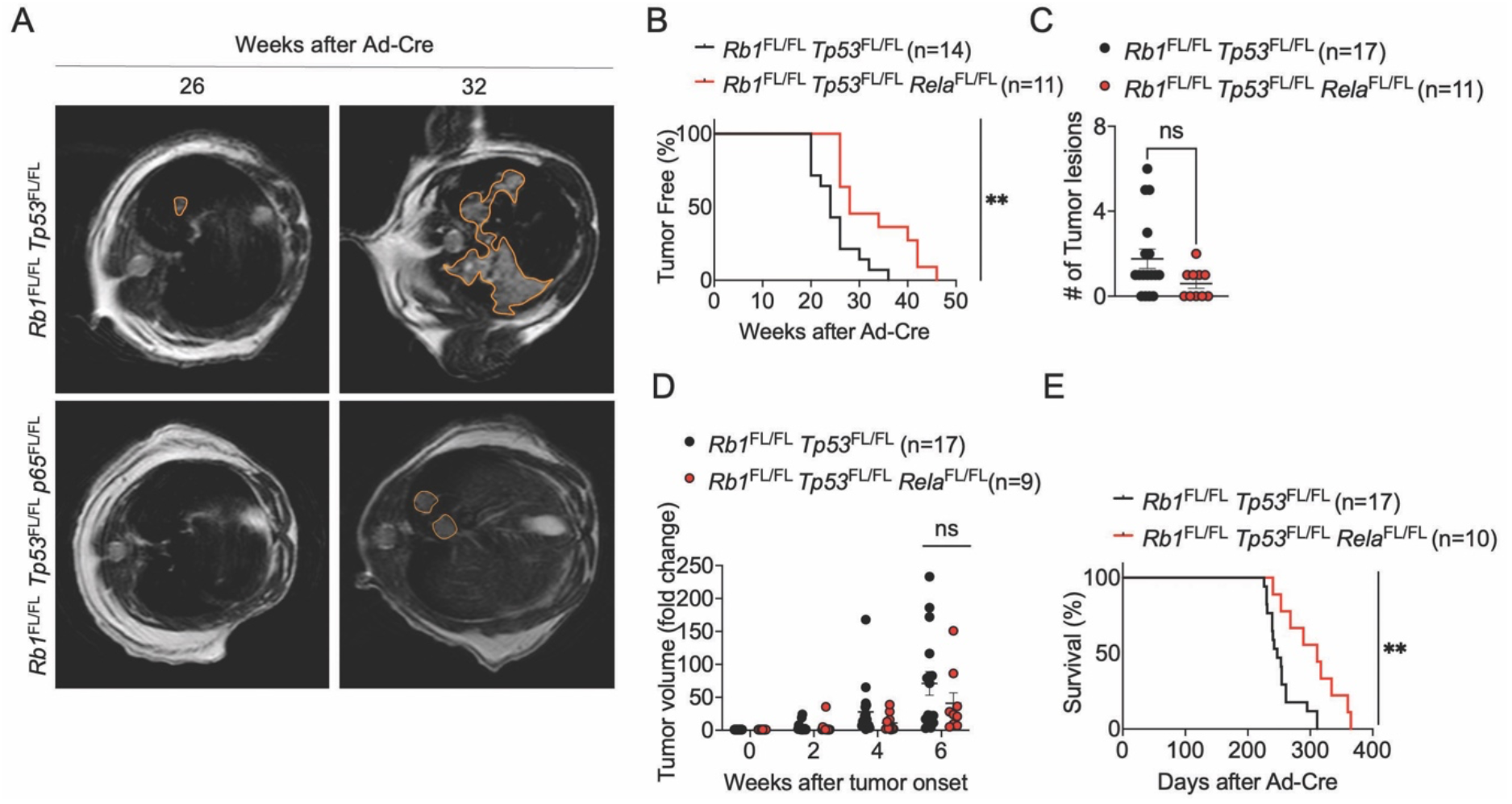
RelA deficiency delays tumor onset and prolongs survival in SCLC. **(A)** Representative images of MRI scans of *Rb1*^FL/FL^ *Tp53*^FL/FL^ (n=17) and *Rb1*^FL/FL^ *Tp53*^FL/FL^ *Rela*^FL/FL^ (n=11) mice 26 and 32 weeks after Ad-Cre inhalation. Tumor areas were marked using Horos Software. **(B)** Graph depicting tumor onset assessed via MR imaging of *Rb1*^FL/FL^ *Tp53*^FL/FL^ and *Rb1*^FL/FL^ *Tp53*^FL/FL^ *Rela*^FL/FL^ mice. **p < 0.005, Log-rank test. **(C)** Graph depicting the number of lung tumor lesions 6 weeks after Ad-Cre inhalation. Each dot represents one mouse. Mean ± SEM are shown. Mann-Whitney Test **(D)** Graph depicting tumor fold change assessed via MR imaging 2-6 weeks after Ad-Cre inhalation for *Rb1*^FL/FL^ *Tp53*^FL/FL^ and *Rb1*^FL/FL^ *Tp53*^FL/FL^ *Rela*^FL/FL^ mice. Each dot represents one mouse. Bars represent mean ± SEM. Manny-Whitney Test. **(E)** Graph depicting survival of mice with indicated genotypes. **p < 0.005, Log-rank test. Data from the same *Rb1*^FL/FL^ *Tp53*^FL/FL^ cohort are included in all figures for comparison.

Macroscopic examination of lung tissues dissected from mice sacrificed at the humane endpoint revealed similar tumor load in *Rb1*^FL/FL^ *Tp53*^FL/FL^ *Rela*^FL/FL^ compared to *Rb1*^FL/FL^ *Tp53*^FL/FL^ mice (Figure 4A). In addition to the lung tumors, SCLC metastasis to the liver was also detected in about half of the mice with no significant difference in metastatic prevalence between the genotypes (Figure 4B). Histopathological analysis of lung sections revealed no difference in tumor morphology, proliferation and immune cell infiltration between *Rb1*^FL/FL^ *Tp53*^FL/FL^ *Rela*^FL/FL^ compared to *Rb1*^FL/FL^ *Tp53*^FL/FL^ mice (Figure 4C, D). RNAseq analysis also failed to reveal considerable gene expression changes, with only 12 genes significantly upregulated and 9 genes significantly downregulated in tumors from *Rb1*^FL/FL^ *Tp53*^FL/FL^ *Rela*^FL/FL^ compared to *Rb1*^FL/FL^ *Tp53*^FL/FL^ mice (Figure 4E, F). Therefore, analysis of tumors at the time of humane sacrifice did not reveal differences in tumor size, proliferation or immune cell infiltration between *Rb1*^FL/FL^ *Tp53*^FL/FL^ *Rela^FL/FL^* and *Rb1*^FL/FL^ *Tp53*^FL/FL^ mice.

**Figure 4.**
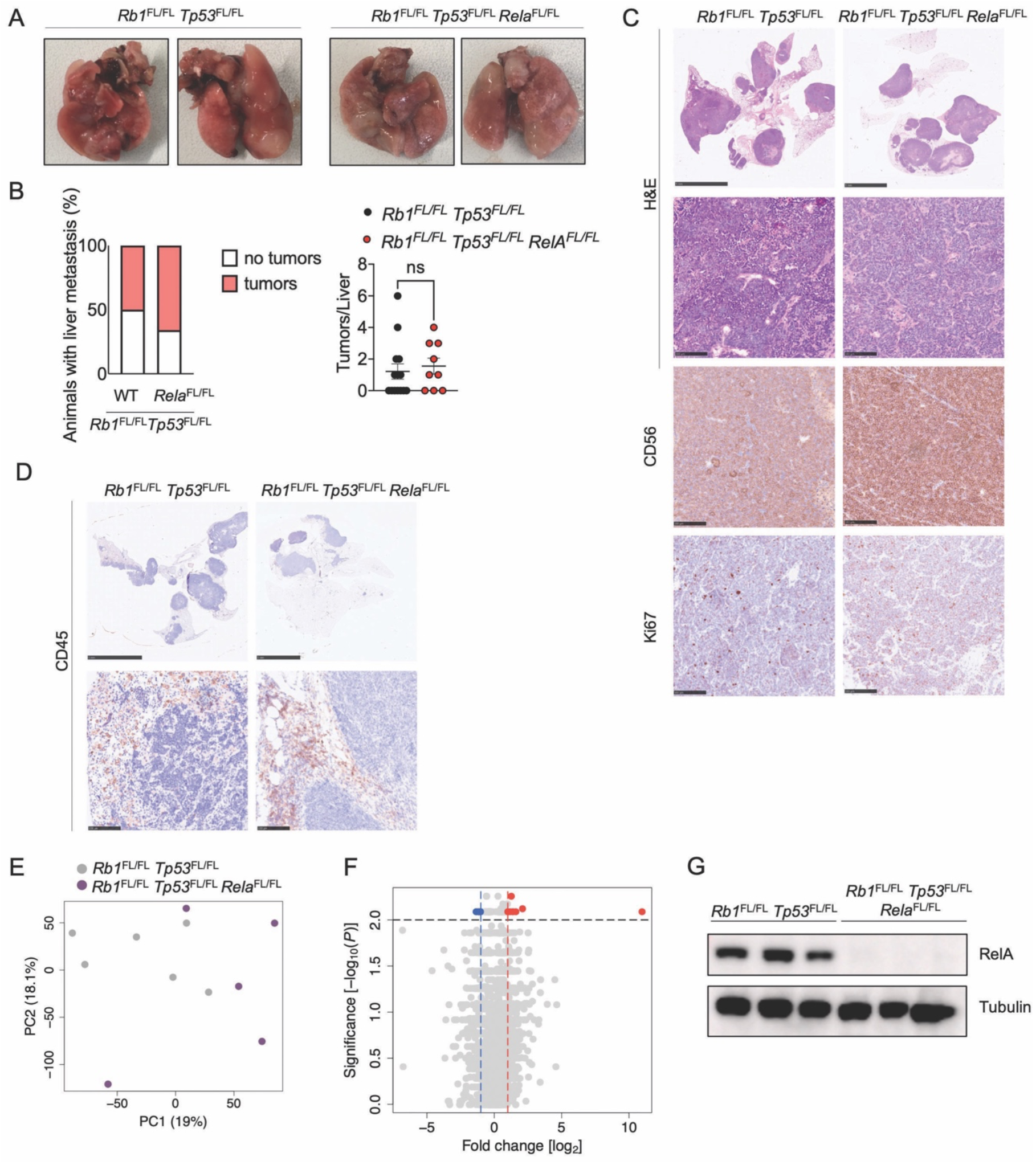
Histological and molecular analysis of lung tumors in *Rb1*^FL/FL^ *Tp53*^FL/FL^ *Rela*^FL/FL^ mice. **(A)** Representative photographs of lungs from *Rb1*^FL/FL^ *Tp53*^FL/FL^ (n=17) and *Rb1*^FL/FL^ *Tp53*^FL/FL^ *Rela*^FL/FL^ (n=10) mice sacrificed at the humane endpoint. **(B)** Graphs depicting the percentage of mice with liver metastasis and the amount of liver tumor lesions at the humane endpoint. n= 9 for *Rb1*^FL/FL^ *Tp53*^FL/FL^ *Rela*^FL/FL^, n=17 for *Rb1*^FL/FL^ *Tp53*^FL/FL^. Mann Whitney test, mean ± SEM. **(C)** Representative images of lung sections from mice with indicated genotypes at the humane endpoint stained with H&E or immunostained for CD56 and for Ki67. Scale bars = 5mm & 100μm (CD56: n=9 for *Rb1*^FL/FL^ *Tp53*^FL/FL^ *Rela*^FL/FL^, n=11 for *Rb1*^FL/FL^ *Tp53*^FL/FL^; Ki67: n=11 for *Rb1*^FL/FL^ *Tp53*^FL/FL^ *Rela*^FL/FL^, n=13 for *Rb1*^FL/FL^ *Tp53*^FL/FL^). **(D)** Representative images of lung sections from mice at the humane endpoint immunostained for CD45. Scale bars = 5 mm (top) and 100μm (bottom), (n=10 for *Rb1*^FL/FL^ *Tp53*^FL/FL^ *Rela*^FL/FL^, n=11 for *Rb1*^FL/FL^ *Tp53*^FL/FL^). **(E-F)** Principal component analysis (PCA) (**E**) and Volcano plot (**F**) of RNA seq data from tumor tissues of *Rb1*^FL/FL^ *Tp53*^FL/FL^ *Rela*^FL/FL^ (n=5) compared to *Rb1*^FL/FL^ *Tp53*^FL/FL^ (n=6) mice. Genes that were found significantly upregulated (p < 0.01, log2(|FC|) > 2) or downregulated (p < 0.01, log2(|FC|) < 2) in tumors from *Rb1*^FL/FL^ *Tp53*^FL/FL^ *Rela*^FL/FL^ (n=5) compared to *Rb1*^FL/FL^ *Tp53*^FL/FL^ (n=6) mice are indicated with red or blue dots, respectively**. (G)** Representative immunoblot analysis of protein extracts from SCLC cell lines isolated from *Rb1*^FL/FL^ *Tp53*^FL/FL^ and *Rb1*^FL/FL^ *Tp53*^FL/FL^ *Rela*^FL/FL^ mice at the humane endpoint (n=3). Data from the same *Rb1*^FL/FL^ *Tp53*^FL/FL^ cohort are included in all figures for comparison.

Immunoblot analysis of tumor cell lines isolated from *Rb1*^FL/FL^ *Tp53*^FL/FL^ *Rela*^FL/FL^ animals showed lack of RelA expression in all samples analyzed (n=3, Figure 4G), demonstrating that RelA was efficiently deleted in the tumors from these mice. Therefore, in contrast to NEMO that appears to be essential for SCLC, RelA promotes tumor initiation and growth but is not necessary for SCLC development. Collectively, these results showed that inhibition of NF-kB signaling by RelA ablation delayed tumor onset and prolonged mouse survival, revealing a tumor-promoting role of NF-kB in SCLC.

### Constitutive NF-κB signaling does not affect SCLC development

Our studies described above showed that inhibition of IKK/NF-kB signaling considerably delayed the onset and progression of SCLC. As a complementary approach, we aimed to assess how constitutively increased activation of IKK/NF-kB signaling might affect SCLC development. For this reason, we employed a mouse model allowing the Cre-mediated expression of a constitutively active IKK2 (IKK2ca) transgene from the ubiquitously expressed Rosa26 locus (*R26^LSL.IKK2ca^* mice) ^39^. We therefore generated *Rb1*^FL/FL^ *Tp53*^FL/FL^ *R26^LSL.IKK2ca^* mice and induced SCLC development by inhalation of Ad-Cre as described above. Based on our findings that NEMO but also RelA ablation delayed tumor development, we hypothesized that persistently elevated NF-kB activation might accelerate and aggravate SCLC. Surprisingly however, MRI-assisted assessment of lung tumor load did not reveal considerable differences in tumor onset or growth between *Rb1*^FL/FL^ *Tp53*^FL/FL^ *R26^LSL.IKK2ca^* and *Rb1*^FL/FL^ *Tp53*^FL/FL^ mice (Figure 5A-D). Consistently, *Rb1*^FL/FL^ *Tp53*^FL/FL^ *R26^LSL.IKK2ca^* mice showed similar overall survival compared to *Rb1*^FL/FL^ *Tp53*^FL/FL^ animals (Figure 5E).

**Figure 5.**
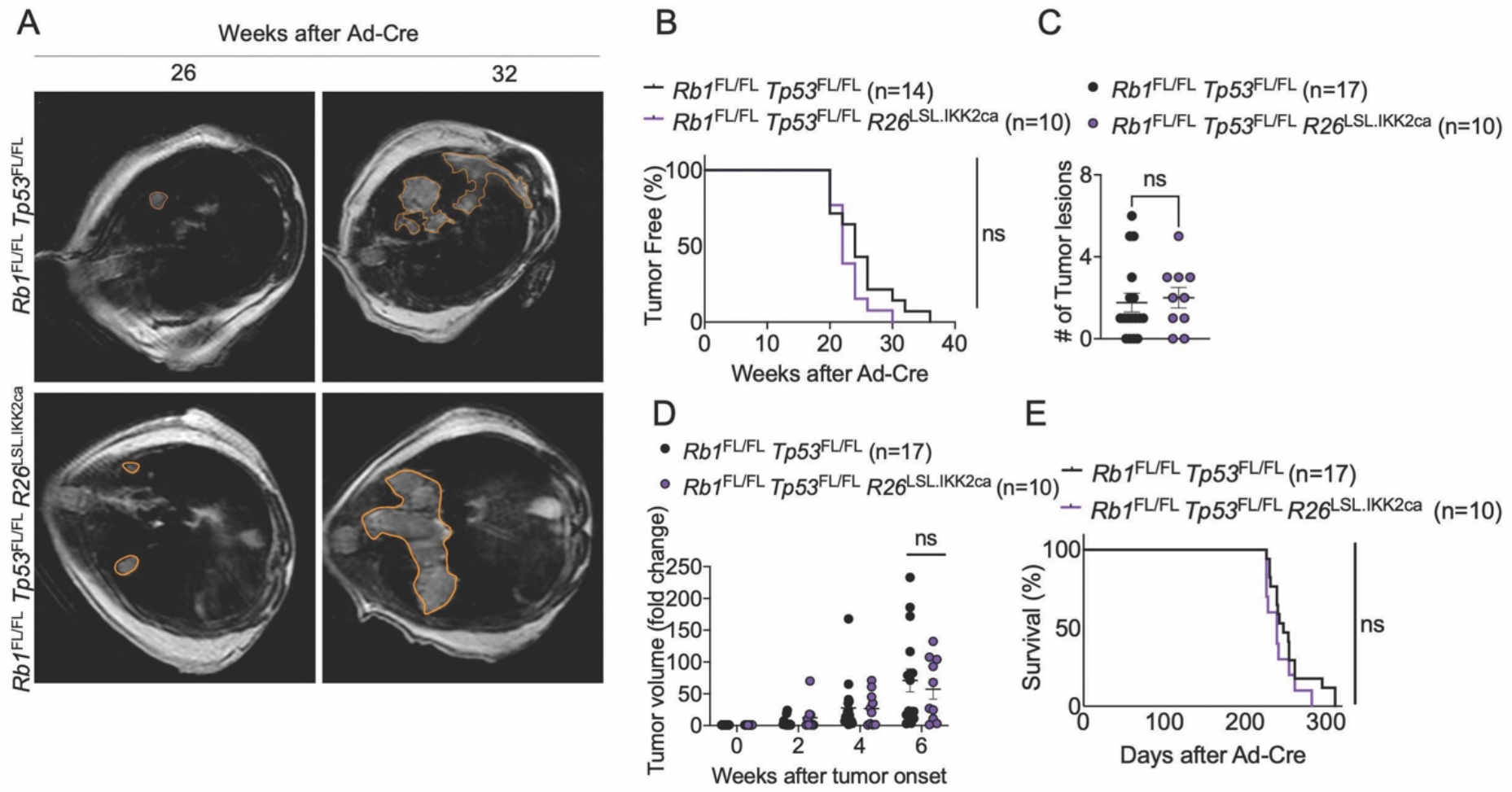
Constitutively increased NF-κB activation did not affect the onset and progression of SCLC. **(A)** Representative images of MRI scans of *Rb1*^FL/FL^ *Tp53*^FL/FL^ (n=17) and *Rb1*^FL/FL^ *Tp53*^FL/FL^ *R26*^LSL.IKK2ca^ (n= 10) mice 26 and 32 weeks after Ad-Cre inhalation. Tumor areas are marked. **(B)** Graph depicting tumor onset assessed via MR imaging of mice with indicated genotypes. **(C)** Graph depicting the number of lung tumor lesions 6 weeks after Ad-Cre inhalation. Each dot represents one mouse. Mean ± SEM are shown. Mann-Whitney Test **(D)** Graph depicting tumor fold change assessed via MR imaging 2-6 weeks after Ad-Cre inhalation for indicated genotypes. Every dot represents one mouse. Bars represent mean ± SEM. Mann-Whitney Test. **(E)** Graph depicting survival of *Rb1*^FL/FL^ *Tp53*^FL/FL^ and *Rb1*^FL/FL^ *Tp53*^FL/FL^ *R26*^LSL.IKK2ca^ mice. Data from the same *Rb1*^FL/FL^ *Tp53*^FL/FL^ cohort are included in all figures for comparison.

Macroscopic examination of dissected lungs from mice sacrificed at the humane endpoint of the experimental protocol revealed similar tumor load in *Rb1*^FL/FL^ *Tp53*^FL/FL^ *R26^LSL.IKK2ca^* and *Rb1*^FL/FL^ *Tp53*^FL/FL^ mice (Figure 6A). Examination of livers from these mice revealed metastasis to this tissue in 2 out of 9 *Rb1*^FL/FL^ *Tp53*^FL/FL^ *R26^LSL.IKK2ca^* mice compared to 7 out 14 *Rb1*^FL/FL^ *Tp53*^FL/FL^ mice, suggesting that expression of IKK2ca could negatively affect the metastatic potential of SCLC (Figure 6B). Moreover, immunohistochemical examination of lung tissues revealed similar tumor burden and morphology in the two genotypes (Figure 6C). Immunostaining for Ki-67 showed similar numbers of proliferating tumor cells in *Rb1*^FL/FL^ *Tp53*^FL/FL^ *R26^LSL.IKK2ca^* compared to *Rb1*^FL/FL^ *Tp53*^FL/FL^ mice (Figure 6C). Furthermore, immunostaining for CD45 revealed no differences in immune cell infiltration in *Rb1*^FL/FL^ *Tp53*^FL/FL^ *R26^LSL.IKK2ca^* compared to *Rb1*^FL/FL^ *Tp53*^FL/FL^ mice, with immune cells surrounding the lesions but generally not found within the tumor mass (Figure 6D). RNAseq analysis revealed considerable changes in gene expression in tumors expressing IKK2ca (Figures 6E-G). Specifically, we found that 152 genes were significantly upregulated in tumors from *Rb1*^FL/FL^ *Tp53*^FL/FL^ *R26^LSL.IKK2ca^* mice compared to *Rb1*^FL/FL^ *Tp53*^FL/FL^ mice, with genes described under the hallmarks “INFLAMMATORY RESPONSE” and “TNFA_SIGNALLING_VIA_NFKB” being significantly enriched within the upregulated gene set (Figures 6E-G). Thus, IKK2ca expression induced the transcriptional upregulation of NF-kB dependent inflammatory genes in SCLC. It is intriguing that the increased expression of inflammatory genes did not enhance immune cell infiltration into the tumors in *Rb1*^FL/FL^ *Tp53*^FL/FL^ *R26^LSL.IKK2ca^* compared to *Rb1*^FL/FL^ *Tp53*^FL/FL^ mice (Figure 6D). Consistent with the elevated expression of NF-kB target genes, immunoblot analysis of IKK2 protein levels revealed strongly increased IKK2ca expression in SCLC tissue isolated from the lungs of *Rb1*^FL/FL^ *Tp53*^FL/FL^ *R26^LSL.IKK2ca^* mice (Figure 6G).

**Figure 6.**
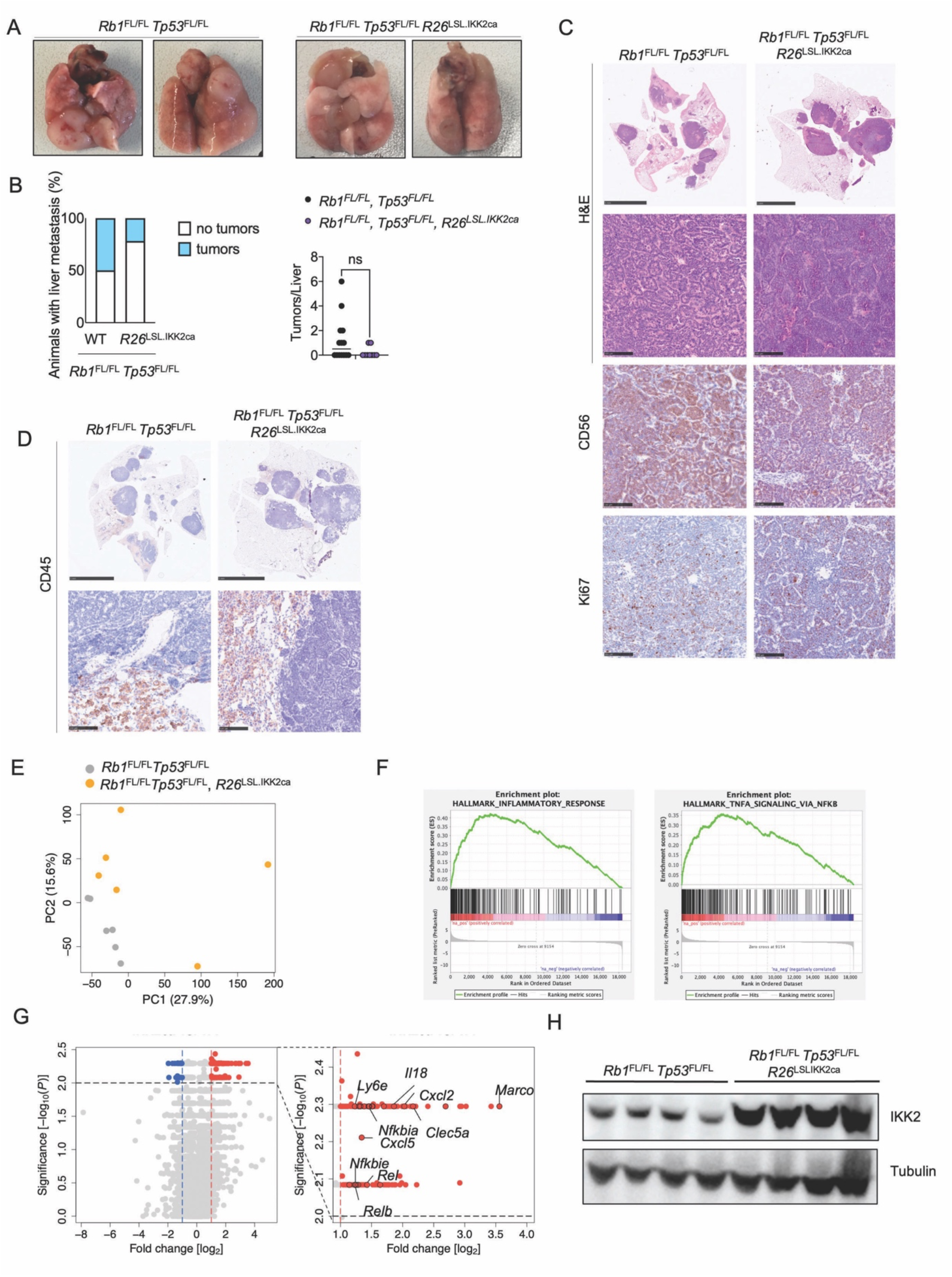
IKK2ca expression increased inflammatory gene transcription but did not alter the morphology or immune cell infiltration of SCLC. **(A)** Representative photographs of lungs from *Rb1*^FL/FL^ *Tp53*^FL/FL^ (n=17) and *Rb1*^FL/FL^ *Tp53*^FL/FL^ *R26*^LSL.IKK2ca^ (n=10) mice sacrificed at the humane endpoint. **(B)** Graphs depicting the percentage of mice with liver metastasis and the amount of liver tumor lesions at the humane endpoint n=9 for *Rb1*^FL/FL^ *Tp53*^FL/FL^ *R26*^LSL.IKK2ca^, n=14 for *Rb1*^FL/FL^ *Tp53*^FL/FL^. Mann-Whitney test, mean ± SEM. **(C)** Representative images of lung sections from mice at the humane endpoint stained with H&E or immunostained for CD56 and for Ki67. Scale bars = 5mm (top) and 100μm (bottom). Samples were taken at the humane endpoint (CD56: n=8 for *Rb1*^FL/FL^ *Tp53*^FL/FL^ *R26*^LSL.IKK2ca^, n=11 for *Rb1*^FL/FL^ *Tp53*^FL/FL^; Ki67: n=7 for *Rb1*^FL/FL^ *Tp53*^FL/FL^ *R26*^LSL.IKK2ca^, n=13 for *Rb1*^FL/FL^ *Tp53*^FL/FL^). **(D)** Representative images of lung sections from *Rb1*^FL/FL^ *Tp53*^FL/FL^ *R26*^LSL.IKK2ca^ (n=8) and *Rb1*^FL/FL^ *Tp53*^FL/FL^ (n=11) mice at the humane endpoint immunostained for CD45. Scale bars = 5mm (top) and 100μm (bottom). **(E-G)** Graphs depicting PCA (**E**), Gene set enrichment analysis (**F**) and Volcano plot (**G**) of RNA seq data from tumor tissues of *Rb1*^FL/FL^ *Tp53*^FL/FL^ *R26*^LSL.IKK2ca^ (n=6) compared to *Rb1*^FL/FL^ *Tp53*^FL/FL^ (n=6) mice. In (**G**), genes that were found significantly upregulated (p < 0.01, log2(|FC|) > 2) or downregulated (p < 0.01, log_2_(|FC|) < 2) in tumors from *Rb1*^FL/FL^ *Tp53*^FL/FL^ *R26*^LSL.IKK2ca^ compared to *Rb1*^FL/FL^ *Tp53*^FL/FL^ mice are indicated with red or blue dots, respectively. **(H)** Representative immunoblot analysis with the indicated antibodies of protein extracts from tumor tissue derived from RP-mice with indicated genotypes at the humane endpoint. Data from the same *Rb1*^FL/FL^ *Tp53*^FL/FL^ cohort are included in all figures for comparison.

Taken together, these results showed that IKK2ca expression caused persistent activation of NF-kB and the transcriptional upregulation of NF-kB target genes. However, this elevated NF-kB activity did not considerably impact on tumor initiation, growth and progression and did not alter the tumor immune landscape in this mouse model of SCLC.

### SCLC development is independent of TNFR1 signaling

TNF signaling via TNFR1 has important functions in tumorigenesis ^12, 49^. However, the role of TNFR1 in SCLC has not been studied and remains unknown. Here, we aimed to address whether TNFR1 signaling contributes to SCLC development using two distinct approaches. One the one hand, we employed TNFR1-deficient (*Tnfr1^-/-^*) mice to assess the role of TNFR1 in both the tumor cells and the cells of the microenvironment. In parallel, we used *Tnfr1*^FL/FL^ mice allowing to assess the tumor cell-intrinsic role of TNFR1. Specifically, we generated *Rb1*^FL/FL^ *Tp53*^FL/FL^ *Tnfr1^-/-^* and *Rb1*^FL/FL^ *Tp53*^FL/FL^ *Tnfr1*^FL/FL^ mice and examined SCLC development after inhalation with Ad-Cre. MRI-assisted assessment of lung tumors revealed that neither tumor cell intrinsic nor systemic TNFR1 deficiency considerably affected the onset and growth of SCLC (Figure 7A-D). Moreover, *Rb1*^FL/FL^ *Tp53*^FL/FL^ *Tnfr1*^-/-^ and *Rb1*^FL/FL^ *Tp53*^FL/FL^ *Tnfr1*^FL/FL^ mice showed similar overall survival compared to *Rb1*^FL/FL^ *Tp53*^FL/FL^ animals (Figure 7E). Therefore, TNFR1 deficiency did not considerably affect tumor initiation, progression and overall survival in SCLC.

**Figure 7.**
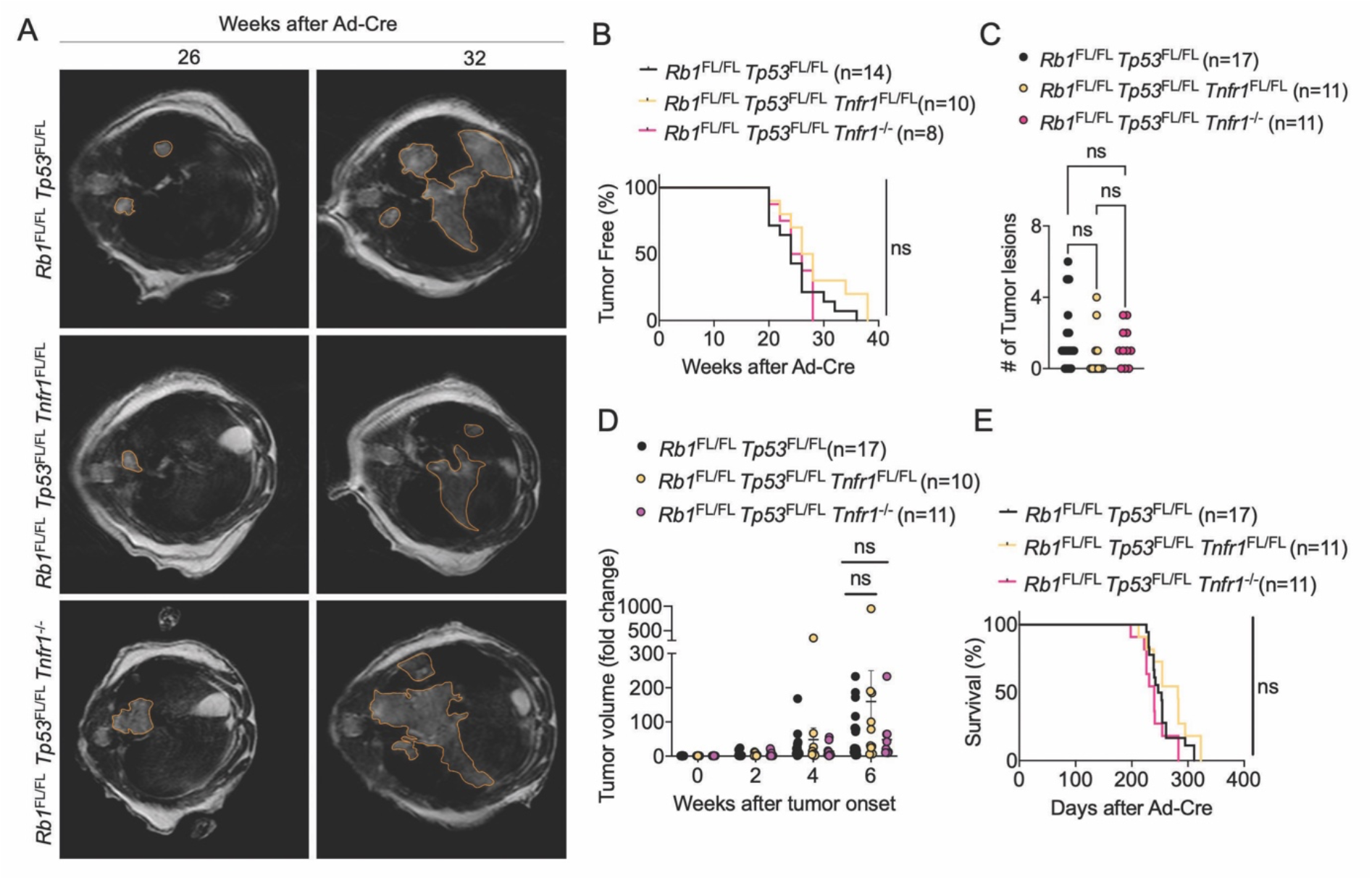
TNFR1 deficiency did not alter SCLC onset or progression. **(A)** Representative images of MRI scans of *Rb1*^FL/FL^ *Tp53*^FL/FL^ (n=17), *Rb1*^FL/FL^ *Tp53*^FL/FL^ *Tnfr1^FL/FL^* (n= 10) and *Rb1*^FL/FL^ *Tp53*^FL/FL^ *Tnfr1^-/-^* (n= 11) mice 26 and 32 weeks after Ad-Cre inhalation. Tumor areas are marked. **(B)** Graph depicting tumor onset assessed via MR imaging of mice with indicated genotypes. **(C)** Graph depicting the number of lung tumor lesions 6 weeks after Ad-Cre inhalation. Each dot represents one mouse. Mean ± SEM are shown. Mann-Whitney Test. **(D)** Graph depicting tumor fold change assessed via MR imaging 2-6 weeks after AdenoCre inhalation for indicated genotypes. Every dot represents one mouse. Bars represent mean ± SEM. Mann-Whitney Test. **(E)** Graph depicting survival of *Rb1*^FL/FL^ *Tp53*^FL/FL^, *Rb1*^FL/FL^ *Tp53*^FL/FL^ *Tnfr1*^FL/FL^ and *Rb1*^FL/FL^ *Tp53*^FL/FL^ *Tnfr1^-/-^* mice. Data from the same *Rb1*^FL/FL^ *Tp53*^FL/FL^ cohort are included in all figures for comparison.

Macroscopic examination of lungs from mice sacrificed at the humane endpoint revealed a similar tumor burden in *Rb1*^FL/FL^ *Tp53*^FL/FL^ *Tnfr1^-/-^* and *Rb1*^FL/FL^ *Tp53*^FL/FL^ *Tnfr1*^FL/FL^ mice compared to *Rb1*^FL/FL^ *Tp53*^FL/FL^ animals (Figure 8A). In addition to the lung tumors, SCLC metastasis to the liver was also detected in several mice with no difference in metastatic prevalence between the genotypes (Figure 8B). Immunohistochemical analysis of lung sections revealed that neither tumor-intrinsic nor ubiquitous lack of TNFR1 could alter tumor morphology and proliferation (Figure 8C). Moreover, immunostaining for CD45 failed to reveal differences in immune cell infiltration between the three genotypes, arguing that inhibition of TNFR1 signaling did not change the immune landscape of SCLC (Figure 8D). Similarly, RNAseq analysis of gene expression in tumors dissected at the humane endpoint did not reveal considerably changed transcription profiles between *Rb1*^FL/FL^ *Tp53*^FL/FL^ *Tnfr1^-/-^* or *Rb1*^FL/FL^ *Tp53*^FL/FL^ *Tnfr1*^FL/FL^ and *Rb1*^FL/FL^ *Tp53*^FL/FL^ mice (Figure 8E,F). Taken together, our results revealed that TNFR1 deficiency, either cell intrinsic or systemic, did not considerably alter SCLC development, arguing that TNFR1 does not play an important role in this type of cancer.

**Figure 8.**
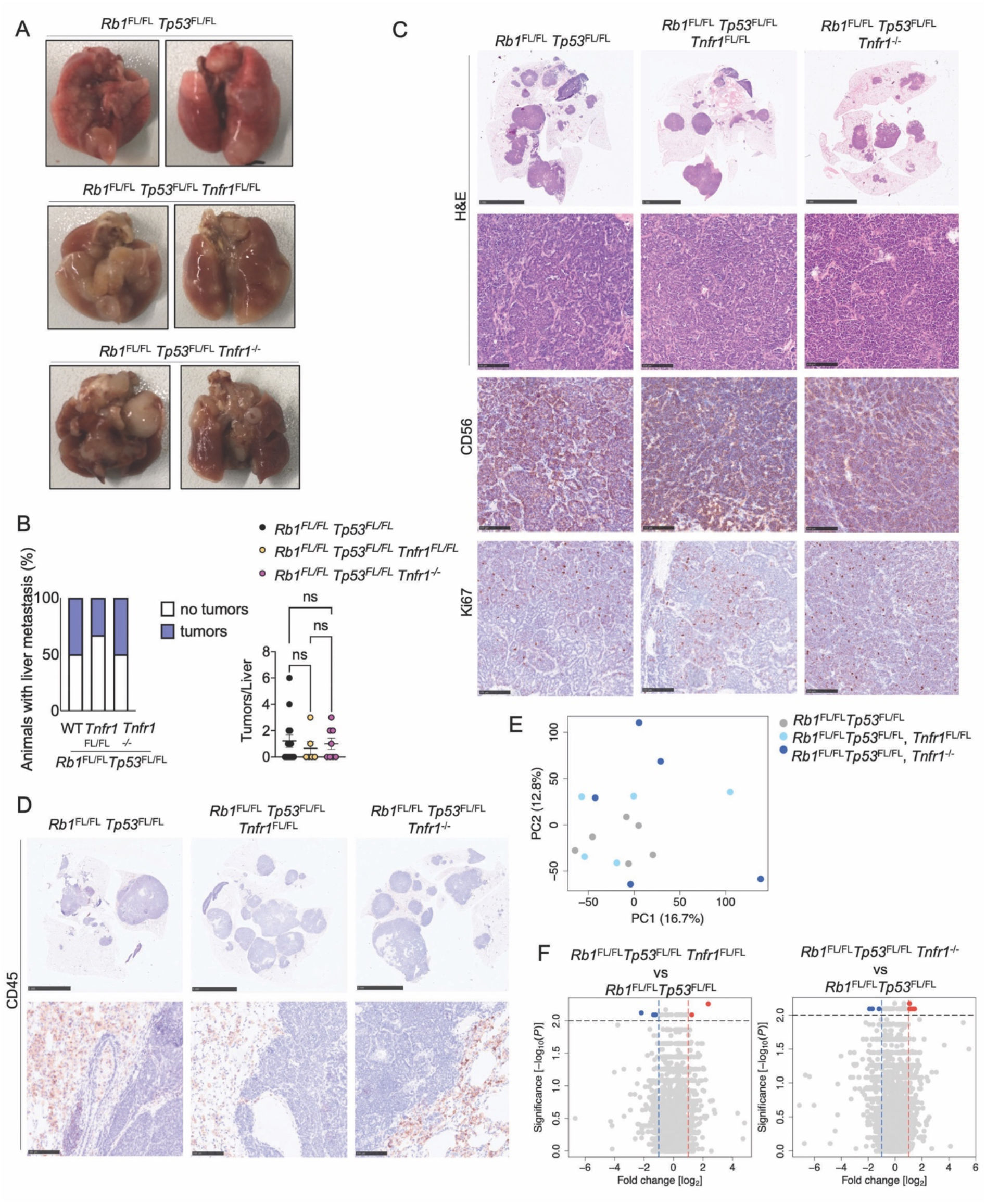
TNFR1 deficiency did not alter SCLC morphology and immune cell infiltration. **(A)** Photographs of lungs of *Rb1*^FL/FL^ *Tp53*^FL/FL^ (n=17), *Rb1*^FL/FL^ *Tp53*^FL/FL^ *Tnfr1^FL/FL^* (n=11) and *Rb1*^FL/FL^ *Tp53*^FL/FL^ *Tnfr1^-/-^* (n=11) mice at the humane endpoint. **(B)** Graphs depicting the percentage of mice with liver metastasis and the amount of liver tumor lesions at the humane endpoint n= 6 for *Rb1*^FL/FL^ *Tp53*^FL/FL^ *Tnfr1*^FL/FL^, n= 8 for *Rb1*^FL/FL^ *Tp53*^FL/FL^ *Tnfr1^-/-^*, n=14 for *Rb1*^FL/FL^ *Tp53*^FL/FL^. Kruskal-Wallis test. **(C)** Representative images of lung sections from mice at the humane endpoint stained with H&E (Scale bars= 5 mm (top) and 100μm (bottom)) or immunostained for CD56 or Ki67 (Scale bars = 100μm) (CD56: n=5 for *Rb1*^FL/FL^ *Tp53*^FL/FL^ *Tnfr1*^FL/FL^, n=8 for *Rb1*^FL/FL^ *Tp53*^FL/FL^ *Tnfr1^-/-^*, n=11 for *Rb1*^FL/FL^ *Tp53*^FL/FL^; Ki67: n=8 for *Rb1*^FL/FL^ *Tp53*^FL/FL^ *Tnfr1*^FL/FL^, n=9 for *Rb1*^FL/FL^ *Tp53*^FL/FL^ *Tnfr1^-/-^*; n=13 for *Rb1*^FL/FL^ *Tp53*^FL/FL^). **(D)** Representative images of lung sections immunostained for CD45. Scale bars = 5mm (top) and 100μm (bottom). Samples were taken at the humane endpoint (n=5 for *Rb1*^FL/FL^ *Tp53*^FL/FL^ *Tnfr1*^FL/FL^, n=8 for *Rb1*^FL/FL^ *Tp53*^FL/FL^ *Tnfr1^-/-^*, n=11 for *Rb1*^FL/FL^ *Tp53*^FL/FL^). **(E-F)** PCA (**E**) and Volcano plot (**F**) of RNA seq data from tumor tissues of *Rb1*^FL/FL^ *Tp53*^FL/FL^ *Tnfr1*^FL/FL^ (n=5) and *Rb1*^FL/FL^ *Tp53*^FL/FL^ *Tnfr1^-/-^* (n=5) mice compared to tumors from *Rb1*^FL/FL^ *Tp53*^FL/FL^ (n=6) mice. Genes that were found significantly upregulated (p < 0.01, log2(|FC|) > 2) or downregulated (p < 0.01, log2(|FC|) < 2) in these comparisons are indicated with red or blue dots, respectively. Data from the same *Rb1*^FL/FL^ *Tp53*^FL/FL^ cohort are included in all figures for comparison.

## Discussion

Despite the progress made in the treatment of different cancer entities during the last decade, SCLC remains a type of cancer with very limited therapeutic options and exceptionally poor prognosis ^1^. Whereas the introduction of immunotherapies had some beneficial effects in patients with SCLC, these were limited to a small fraction (about 15%) of the patients ^1^. Therefore, new therapeutic targets for SCLC are urgently needed. Here, we have addressed the role of TNFR1 and NF-κB signaling in development and progression of SCLC in a well-established and relevant mouse model of the disease. Surprisingly, we found that TNFR1 deficiency, either tumor cell intrinsic or systemic, did not affect SCLC development, in contrast to the important role of TNF signaling in other types of cancer ^12, 49^. This finding suggests that, at least in this specific mouse model driven by acute inactivation of the two important tumor suppressors RB1 and TP53, TNF-mediated inflammatory and cell death signaling is not critically involved. It should be noted however, that these findings do not exclude a role of TNF in human patients with SCLC associated with comorbidities such as chronic obstructive pulmonary disease, where lung inflammation may contribute to tumor progression.

Our results revealed a critical role of IKK/NF-kB signaling in the development of SCLC. Ablation of NEMO significantly delayed tumor onset, slowed tumor growth and considerably prolonged survival in mice with SCLC induced by combined inactivation of RB1 and TP53. Interestingly, we found that SCLC cell lines isolated from *Rb1*^FL/FL^ *Tp53*^FL/FL^ *Nemo^FL/FL^* mice displayed expression of NEMO, showing that these were derived from cells that failed to recombine the NEMO floxed allele. Considering the critical pro-survival function of NEMO, these results suggest that NEMO deficiency causes the death of transformed cells early on resulting in strong counter-selection of these cells, with the tumors eventually developing in these animals arising from rare cells that underwent Cre-mediated recombination of the *Rb1*^FL/FL^ *Tp53*^FL/FL^ but not of the *Nemo^FL/FL^* alleles. NEMO exhibits both NF-kB-dependent and NF-kB-independent pro-survival functions ^31, 48^. Our findings that RelA ablation delayed the onset and growth of SCLC resulting in considerably prolonged mouse survival provided experimental evidence that RelA-dependent NF-kB-dependent gene transcription also plays an important role in SCLC. RelA deficiency had a less pronounced effect compared to loss of NEMO, suggesting that NEMO ablation suppresses SCLC development by both NF-kB-dependent and -independent mechanisms. Importantly, analysis of SCLC cell lines isolated from lungs of Ad-Cre-inhaled *Rb1*^FL/FL^ *Tp53*^FL/FL^ *Rela^FL/FL^* mice sacrificed at the humane endpoint revealed efficient ablation of RelA, showing that RelA-deficient cells can give rise to SCLC, in contrast to loss of NEMO that appears to be incompatible with SCLC development. Together, these studies suggest a dual function of the IKK/NF-kB signaling pathway in SCLC. Complete inhibition of canonical IKK/NF-kB signaling by NEMO deficiency prevented the development of SCLC, most likely by sensitizing RB1-TP53 double-deficient cells to death, whereas RelA knockout had a less pronounced effect in delaying tumor development and progression. Interestingly, expression of constitutively active IKK2ca did not exacerbate SCLC development, showing that persistently elevated IKK/NF-kB activity by did not provide an advantage to the tumors. Surprisingly, despite the upregulation of inflammatory gene expression, *Rb1*^FL/FL^ *Tp53*^FL/FL^ *R26^LSL.IKK2ca^* mice did not show increased infiltration of immune cells within the tumor mass. Taken together, our results revealed a tumor-promoting role of IKK/NF-kB signaling in SCLC, in line with previous studies showing that NF-kB critically contributes to Kras mutation-driven lung adenocarcinoma ^25, 26, 27^. These findings suggest that IKK/NF-kB signaling could provide a promising therapeutic target in SCLC and warrant further studies experimentally assessing the effect of NF-kB pathway inhibition on established SCLC.

## Acknowledgements

We thank A. Florin, F. Schneider, J. Kuth, C. Uthoff-Hachenberg, E. Stade and E. Gareus for excellent technical assistance, as well as C.M. Bebber, A. Androulidaki and H. Schuenke for help with the *Rb1*^FL/FL^*Tp53*^FL/FL^ mouse model. We also thank H. Grüll for support with MRI imaging, H.C. Reinhardt and R. Thomas for valuable discussions and G. Kollias for providing *Tnfr1*^FL/FL^ mice. This work was supported by funding from the Deutsche Forschungsgemeinschaft (DFG, German Research Foundation), project SFB1399 (Project No. 413326622), to M. Pasparakis, M. Peifer and R. Buettner, and from the Federal Ministry of Education and Research (BMBF, e:med project InCa, Grant No. 01ZX1901A) to M.P..

## Author contributions

L.K. designed and performed all experiments, analyzed the data and drafted and revised the manuscript. T.-P.Y. and M. Peifer analyzed RNA sequencing data sets. M.S. and R.B. contributed to the histologic analysis. M. Pasparakis designed and supervised the study and wrote the manuscript together with L.K.

## Conflict of interest

R.B. is an employee of Targos Molecular Pathology. The other authors declare no competing interests.

## References

1. Rudin CM, Brambilla E, Faivre-Finn C, Sage J. Small-cell lung cancer. Nat Rev Dis Primers 2021, 7(1): 3.

2. Junker K, Wiethege T, Muller KM. Pathology of small-cell lung cancer. J Cancer Res Clin Oncol 2000, 126(7): 361–368.

3. Rom WN, Hay JG, Lee TC, Jiang Y, Tchou-Wong KM. Molecular and genetic aspects of lung cancer. Am J Respir Crit Care Med 2000, 161 (4 Pt 1): 1355–1367.

4. Wistuba, II, Gazdar AF, Minna JD. Molecular genetics of small cell lung carcinoma. Semin Oncol 2001, 28(2 Suppl 4): 3–13.

5. Sandler AB. Chemotherapy for small cell lung cancer. Semin Oncol 2003, 30(1): 9–25.

6. Harbour JW, Lai SL, Whang-Peng J, Gazdar AF, Minna JD, Kaye FJ. Abnormalities in structure and expression of the human retinoblastoma gene in SCLC. Science 1988, 241(4863): 353–357.

7. George J, Lim JS, Jang SJ, Cun Y, Ozretic L, Kong G, et al. Comprehensive genomic profiles of small cell lung cancer. Nature 2015, 524(7563): 47–53.

8. Peifer M, Fernandez-Cuesta L, Sos ML, George J, Seidel D, Kasper LH, et al. Integrative genome analyses identify key somatic driver mutations of small-cell lung cancer. Nat Genet 2012, 44(10): 1104–1110.

9. Sekido Y, Fong KM, Minna JD. Molecular genetics of lung cancer. Annu Rev Med 2003, 54: 73–87.

10. Gouyer V, Gazzeri S, Bolon I, Drevet C, Brambilla C, Brambilla E. Mechanism of retinoblastoma gene inactivation in the spectrum of neuroendocrine lung tumors. Am J Respir Cell Mol Biol 1998, 18(2): 188–196.

11. Hanahan D, Weinberg RA. Hallmarks of cancer: the next generation. Cell 2011, 144(5): 646–674.

12. Montfort A, Colacios C, Levade T, Andrieu-Abadie N, Meyer N, Segui B. The TNF Paradox in Cancer Progression and Immunotherapy. Front Immunol 2019, 10: 1818.

13. Moore RJ, Owens DM, Stamp G, Arnott C, Burke F, East N, et al. Mice deficient in tumor necrosis factor-alpha are resistant to skin carcinogenesis. Nat Med 1999, 5(7): 828–831.

14. Park EJ, Lee JH, Yu GY, He G, Ali SR, Holzer RG, et al. Dietary and genetic obesity promote liver inflammation and tumorigenesis by enhancing IL-6 and TNF expression. Cell 2010, 140(2): 197–208.

15. Varfolomeev E, Vucic D. Intracellular regulation of TNF activity in health and disease. Cytokine 2018, 101: 26–32.

16. Pasparakis M. Regulation of tissue homeostasis by NF-kappaB signalling: implications for inflammatory diseases. Nat Rev Immunol 2009, 9(11): 778–788.

17. Taniguchi K, Karin M. NF-kappaB, inflammation, immunity and cancer: coming of age. Nat Rev Immunol 2018, 18(5): 309–324.

18. Hayden MS, Ghosh S. NF-kappaB, the first quarter-century: remarkable progress and outstanding questions. Genes Dev 2012, 26(3): 203–234.

19. Karin M. Nuclear factor-kappaB in cancer development and progression. Nature 2006, 441(7092): 431–436.

20. Greten FR, Eckmann L, Greten TF, Park JM, Li ZW, Egan LJ, et al. IKKbeta links inflammation and tumorigenesis in a mouse model of colitis-associated cancer. Cell 2004, 118(3): 285–296.

21. Pratt MA, Tibbo E, Robertson SJ, Jansson D, Hurst K, Perez-Iratxeta C, et al. The canonical NF-kappaB pathway is required for formation of luminal mammary neoplasias and is activated in the mammary progenitor population. Oncogene 2009, 28(30): 2710–2722.

22. Liu M, Sakamaki T, Casimiro MC, Willmarth NE, Quong AA, Ju X, et al. The canonical NF-kappaB pathway governs mammary tumorigenesis in transgenic mice and tumor stem cell expansion. Cancer Res 2010, 70(24): 10464–10473.

23. Connelly L, Barham W, Onishko HM, Sherrill T, Chodosh LA, Blackwell TS, et al. Inhibition of NF-kappa B activity in mammary epithelium increases tumor latency and decreases tumor burden. Oncogene 2011, 30(12): 1402–1412.

24. Kim C, Pasparakis M. Epidermal p65/NF-kappaB signalling is essential for skin carcinogenesis. EMBO Mol Med 2014, 6(7): 970–983.

25. Xia Y, Yeddula N, Leblanc M, Ke E, Zhang Y, Oldfield E, et al. Reduced cell proliferation by IKK2 depletion in a mouse lung-cancer model. Nat Cell Biol 2012, 14(3): 257–265.

26. Basseres DS, Ebbs A, Levantini E, Baldwin AS. Requirement of the NF-kappaB subunit p65/RelA for K-Ras-induced lung tumorigenesis. Cancer Res 2010, 70(9): 3537–3546.

27. Meylan E, Dooley AL, Feldser DM, Shen L, Turk E, Ouyang C, et al. Requirement for NF-kappaB signalling in a mouse model of lung adenocarcinoma. Nature 2009, 462(7269): 104–107.

28. Dajee M, Lazarov M, Zhang JY, Cai T, Green CL, Russell AJ, et al. NF-kappaB blockade and oncogenic Ras trigger invasive human epidermal neoplasia. Nature 2003, 421(6923): 639–643.

29. van Hogerlinden M, Rozell BL, Ahrlund-Richter L, Toftgard R. Squamous cell carcinomas and increased apoptosis in skin with inhibited Rel/nuclear factor-kappaB signaling. Cancer Res 1999, 59(14): 3299–3303.

30. Lind MH, Rozell B, Wallin RP, van Hogerlinden M, Ljunggren HG, Toftgard R, et al. Tumor necrosis factor receptor 1-mediated signaling is required for skin cancer development induced by NF-kappaB inhibition. Proc Natl Acad Sci U S A 2004, 101(14): 4972–4977.

31. Kondylis V, Polykratis A, Ehlken H, Ochoa-Callejero L, Straub BK, Krishna-Subramanian S, et al. NEMO Prevents Steatohepatitis and Hepatocellular Carcinoma by Inhibiting RIPK1 Kinase Activity-Mediated Hepatocyte Apoptosis. Cancer Cell 2015, 28(5): 582–598.

32. Luedde T, Beraza N, Kotsikoris V, van Loo G, Nenci A, De Vos R, et al. Deletion of NEMO/IKKgamma in liver parenchymal cells causes steatohepatitis and hepatocellular carcinoma. Cancer Cell 2007, 11(2): 119–132.

33. Maeda S, Kamata H, Luo JL, Leffert H, Karin M. IKKbeta couples hepatocyte death to cytokine-driven compensatory proliferation that promotes chemical hepatocarcinogenesis. Cell 2005, 121 (7): 977–990.

34. Hopewell EL, Zhao W, Fulp WJ, Bronk CC, Lopez AS, Massengill M, et al. Lung tumor NF-kappaB signaling promotes T cell-mediated immune surveillance. J Clin Invest 2013, 123(6): 2509–2522.

35. Vooijs M, van der Valk M, te Riele H, Berns A. Flp-mediated tissue-specific inactivation of the retinoblastoma tumor suppressor gene in the mouse. Oncogene 1998, 17(1): 1–12.

36. Jonkers J, Meuwissen R, van der Gulden H, Peterse H, van der Valk M, Berns A. Synergistic tumor suppressor activity of BRCA2 and p53 in a conditional mouse model for breast cancer. Nat Genet 2001, 29(4): 418–425.

37. Schmidt-Supprian M, Bloch W, Courtois G, Addicks K, Israel A, Rajewsky K, et al. NEMOlIKK gamma-deficient mice model incontinentia pigmenti. Mol Cell 2000, 5(6): 981–992.

38. Luedde T, Heinrichsdorff J, de Lorenzi R, De Vos R, Roskams T, Pasparakis M. IKK1 and IKK2 cooperate to maintain bile duct integrity in the liver. Proc Natl Acad Sci U S A 2008, 105(28): 9733–9738.

39. Sasaki Y, Derudder E, Hobeika E, Pelanda R, Reth M, Rajewsky K, et al. Canonical NF-kappaB activity, dispensable for B cell development, replaces BAFF-receptor signals and promotes B cell proliferation upon activation. Immunity 2006, 24(6): 729–739.

40. Van Hauwermeiren F, Armaka M, Karagianni N, Kranidioti K, Vandenbroucke RE, Loges S, et al. Safe TNF-based antitumor therapy following p55TNFR reduction in intestinal epithelium. J Clin Invest 2013, 123(6): 2590–2603.

41. Pfeffer K, Matsuyama T, Kundig TM, Wakeham A, Kishihara K, Shahinian A, et al. Mice deficient for the 55 kd tumor necrosis factor receptor are resistant to endotoxic shock, yet succumb to L. monocytogenes infection. Cell 1993, 73(3): 457–467.

42. Bray NL, Pimentel H, Melsted P, Pachter L. Near-optimal probabilistic RNA-seq quantification. Nat Biotechnol 2016, 34(5): 525–527.

43. Pimentel H, Bray NL, Puente S, Melsted P, Pachter L. Differential analysis of RNA-seq incorporating quantification uncertainty. Nat Methods 2017, 14(7): 687–690.

44. Subramanian A, Tamayo P, Mootha VK, Mukherjee S, Ebert BL, Gillette MA, et al. Gene set enrichment analysis: a knowledge-based approach for interpreting genomewide expression profiles. Proc Natl Acad Sci U S A 2005, 102(43): 15545–15550.

45. Meuwissen R, Linn SC, Linnoila RI, Zevenhoven J, Mooi WJ, Berns A. Induction of small cell lung cancer by somatic inactivation of both Trp53 and Rb1 in a conditional mouse model. Cancer Cell 2003, 4(3): 181–189.

46. Kontogianni K, Nicholson AG, Butcher D, Sheppard MN. CD56: a useful tool for the diagnosis of small cell lung carcinomas on biopsies with extensive crush artefact. J Clin Pathol 2005, 58(9): 978–980.

47. Hayden MS, West AP, Ghosh S. NF-kappaB and the immune response. Oncogene 2006, 25(51): 6758–6780.

48. Vlantis K, Wullaert A, Polykratis A, Kondylis V, Dannappel M, Schwarzer R, et al. NEMO Prevents RIP Kinase 1-Mediated Epithelial Cell Death and Chronic Intestinal Inflammation by NF-kappaB-Dependent and -Independent Functions. Immunity 2016, 44(3): 553–567.

49. Balkwill F. Tumour necrosis factor and cancer. Nat Rev Cancer 2009, 9(5): 361–371.

